# Reliability and Generalizability of Similarity-Based Fusion of MEG and fMRI Data in Human Ventral and Dorsal Visual Streams

**DOI:** 10.1101/451526

**Authors:** Yalda Mohsenzadeh, Caitlin Mullin, Benjamin Lahner, Radoslaw Cichy, Aude Oliva

## Abstract

To build a representation of what we see, the human brain recruits regions throughout the visual cortex in cascading sequence. Recently, an approach was proposed to evaluate the dynamics of visual perception in high spatiotemporal resolution at the scale of the whole brain. This method combined functional magnetic resonance imaging (fMRI) data with magnetoencephalography (MEG) data using representational similarity analysis and revealed a hierarchical progression from primary visual cortex through the dorsal and ventral streams. To assess the replicability of this method, here we present results of a visual recognition neuro-imaging fusion experiment, and compare them within and across experimental settings. We evaluated the reliability of this method by assessing the consistency of the results under similar test conditions, showing high agreement within participants. We then generalized these results to a separate group of individuals and visual input by comparing them to the fMRI-MEG fusion data of Cichy et al (2016), revealing a highly similar temporal progression recruiting both the dorsal and ventral streams. Together these results are a testament to the reproducibility of the fMRI-MEG fusion approach and allows for the interpretation of these spatiotemporal dynamic in a broader context.

## 1. Introduction

To solve visual object recognition, the human brain has developed a particular cortical topology within the ventral and dorsal streams, recruiting regions in cascading sequence, to quickly build a representation of what we see (i.e. [1–9]).

To reveal the complex neural dynamics underlying visual object recognition, neural responses must be resolved in both space and time simultaneously [10–15]. Towards this aim, Cichy and collaborators proposed a novel approach to combine functional magnetic resonance imaging (fMRI) with magnetoencephalography (MEG) termed MEG-fMRI fusion [8,9,16–18]. The results revealed the dynamics of the visual processing cascade. Neural responses first emerge in the occipital pole (V1, V2, V3) at around 80 msec, and then progress in the anterior direction along the ventral (i.e. lateral-occipital cortex LO, ventral occipital cortex VO, temporal occipital cortex TO and parahippocampal cortex PHC) and dorsal (intraparietal sulcus regions) visual streams within 110-170 msec after image onset.

The consistency of these results with established findings [19–25] suggests that fMRI-MEG fusion is an appropriate analytical tool to non-invasively evaluate the spatiotemporal mechanisms of perception. To assess the power of the fusion technique to yield replicable results on visual recognition dynamics in the human ventral and dorsal streams [26–28], here we replicate [9]. Specifically, we first evaluated the reliability of the fusion method at capturing the spatiotemporal dynamics of visual perception by assessing the neural agreement of visually similar experiences within individuals, asking: ‘Do similar visual experiences obey similar spatiotemporal stages in the brain?’ Given the known variation across brain regions for separate visual category input (e.g. [1,29–34]), our second objective was to determine the generalizability of these patterns: ‘For a given task, which spatiotemporal properties are reproducible across diverse visual input and independent observer groups?’

## 2. Materials and Methods

This paper presents two independent experiments in which fMRI and MEG data were acquired when observers look at pictures of natural images. The fMRI and MEG data of Experiment 1 are original to this current work. Data of Experiment 2 has been published originally in [9] (Experiment 2).

### 2.1. Participants

Two separate groups of fifteen right-handed volunteers with normal or corrected to normal vision participated in Experiment 1 (9 female, 27.87 ± 5.17 years old) and Experiment 2 (5 female, 26.6 ± 5.18 years old, see [9]). The participants signed an informed consent form and were compensated for their participation. Both studies were conducted in accordance with the Declaration of Helsinki and approved by the Institutional Review Board of Massachusetts Institute of Technology.

### 2.2. Stimulus Set

In Experiment 1, the stimulus set included *twin* sets of 78 real-world natural images each (156 images total). Twin-set 1 and Twin-set 2 each contained an image with the same verbal semantic description (see Figure 1a, i.e. a pair of sunflowers, giraffes, horses, etc.). The sets were not significantly different on a collection of low level image statistics [35,36]. The stimulus set of Experiment 2 consisted of 118 natural images of objects [9] from the ImageNet dataset [37]. In both experiments, participants performed the same orthogonal vigilance task. See Appendix A for details on the Experimental Design.

**Figure 1.**
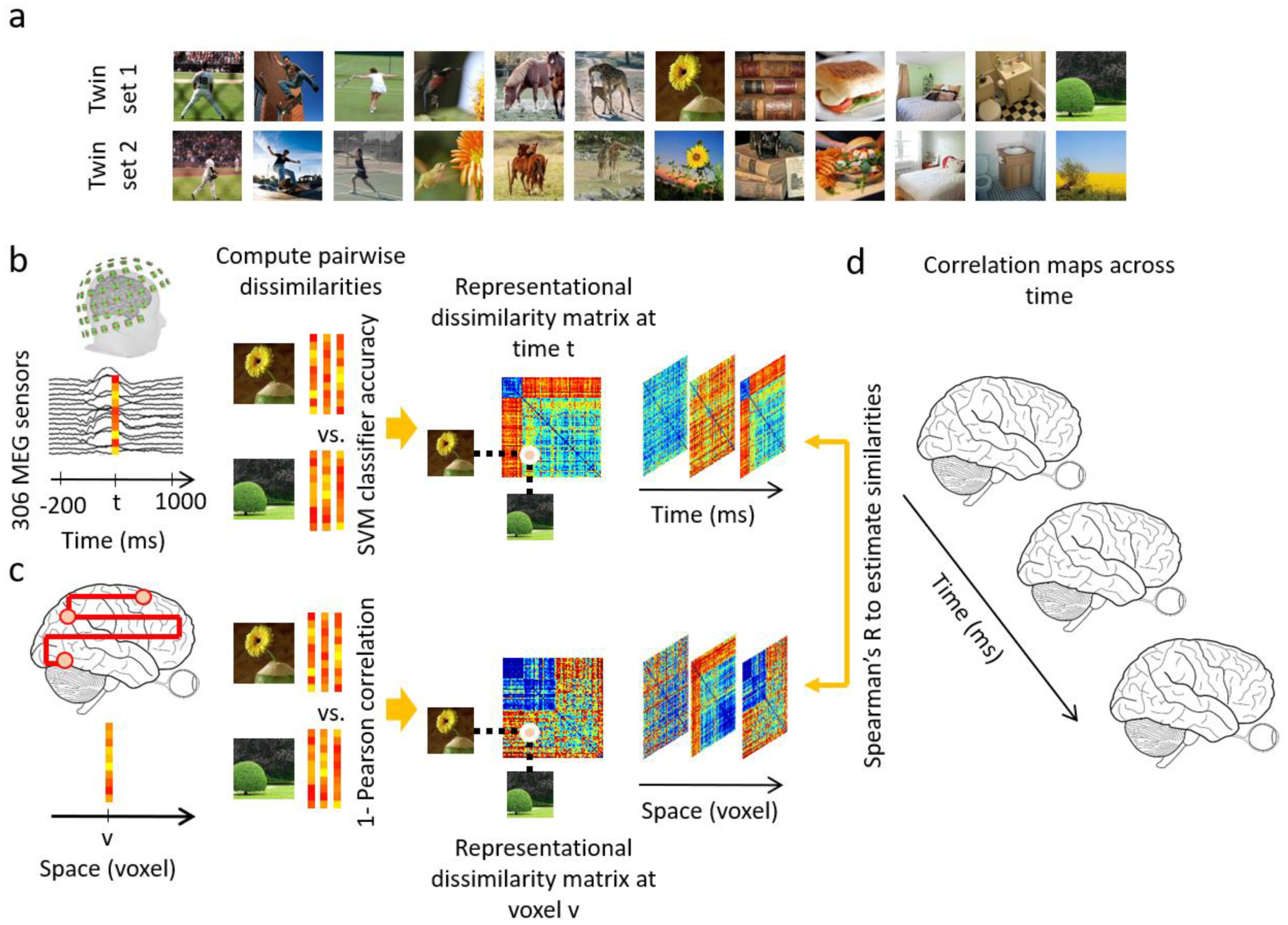
Examples of stimuli of Experiment 1 and analysis scheme: (**a**) Examples of experimental condition: The stimulus set consisted of two sets (termed *twin-sets 1 and 2*) of 78 object images each ordered in pairs, such that each set was of the same semantic content but different in appearance. (**b**) MEG multivariate analysis. MEG data were extracted from −200 msec to 1000 msec with respect to stimulus onset. The 306 dimensional MEG sensor data at each time point t were arranged in vectors. For each pair of conditions, the performance of a SVM classifier in discriminating the conditions based on vector patterns was used as a measure of dissimilarity between the pair of conditions. A representational dissimilarity matrix at each time point t was populated using these pairwise dissimilarities; (**c**) fMRI multivariate analysis. At each voxel v, the activation patterns in a sphere with radius of 4 voxels were extracted and pairwise dissimilarities of conditions were computed (1-Pearson’s R) yielding a representational dissimilarity matrix assigned to the voxel v at the center of the sphere; (**d**) fMRI-MEG similarity based fusion. The MEG RDM representation at each time point t were compared with the fMRI RDM representations at each voxel v by computing Spearman’s R correlations. These computations resulted in correlation maps over the whole brain and across time.

### 2.3. fMRI and MEG Acquisition

MEG and fMRI data for Experiment 1 were acquired in separate sessions, similar to Experiment 2 in [9]. Images were presented 500 msec in all conditions. See Appendix B for data acquisition detail of both experiments.

### 2.4. Data Analyses

We performed several data analyses to test the robustness and generalizability of our results. First, we performed a *full brain fMRI-MEG fusion* [8,9,16,17], which uses representational similarity analysis [38,39] to map MEG and fMRI data into a common space (see Appendix C for details). To summarize, the MEG data were analyzed in a time-resolved manner with 1 msec resolution. MEG sensor data at each time point were arranged in pattern vectors for each stimulus condition and repetition. These vectors were then used to train support vector machines to classify each pair of conditions. The performance of the binary SVM classifiers computed with leave-one-out cross validation procedure were interpreted as a pairwise dissimilarity measure (higher decoding indicates larger dissimilarity) and used to construct a condition by condition representational dissimilarity matrix (RDM) per time point (Figure 1b). The fMRI data were analyzed with a searchlight approach to construct the representational dissimilarity matrices in a voxel-resolved fashion. At every voxel in the brain, the condition-specific voxel patterns in its vicinity were extracted and pairwise condition-specific dissimilarities (1 - Pearson’s R) were computed to create a condition by condition fMRI RDM assigned to that voxel (Figure 1c). Then, in the similarity space of RDMs, MEG and fMRI data were directly compared (Spearman’s R) to integrate high spatial resolution of fMRI with high temporal resolution of MEG. In detail, for a given time point the MEG RDM was correlated with fMRI RDMs specified with the searchlight method resulting in a 3D correlation map. Repeating the procedure for all time points as depicted in Figure 1d resulted in a 4D spatiotemporal correlation map per individual. The correlation maps were averaged over individuals and significant correlations were determined with permutation tests and multiple comparison corrections over time and space with cluster correction method (N=15 in each experiment; cluster-definition threshold of 0.001 and cluster size threshold of 0.01). The detail on statistical tests are presented in Appendix E.

Second, to quantitatively compare the spatiotemporal neural dynamics within and across the experiments and show how reliable they are, we performed two types of anatomically-defined *region-of-interest (ROI) based analysis* along ventral and dorsal streams (see Appendix D for detail): 1) a *spatially restricted searchlight voxel-wise analysis*: in this analysis, the searchlight based MEG-fMRI fusion correlation time series are averaged over voxels within an ROI; 2) the conventional *ROI-based analysis*: in this analysis, the condition-specific voxel responses within an ROI are extracted, and the pairwise condition-specific response dissimilarities are computed to create an ROI RDM which is then compared with time-resolved MEG RDMs resulting in correlation time series (as in [9]).

## 3. Results

### 3.1. Reliability of similarity-based fMRI-MEG fusion method

We first assessed the reliability of the fMRI-MEG similarity-based data fusion method. For this we applied fMRI-MEG fusion separately to the two sets of images making up the Twins-set (i.e. Twinset 1 and Twin-set 2) and compared the results.

Figure 2 and 3 show the spatiotemporal dynamics of visual perception in the ventral and dorsal visual pathways respectively for the Twins-sets over the time course of 1000 msec. Qualitatively examining these results reveal that the neural responses start in occipital pole around 70-90 msec after stimulus onset, followed by neural responses in anterior direction along the ventral stream (Figure 2ac), and across the dorsal stream up to the inferior parietal cortex (Figure 3ac).

**Figure 2.**
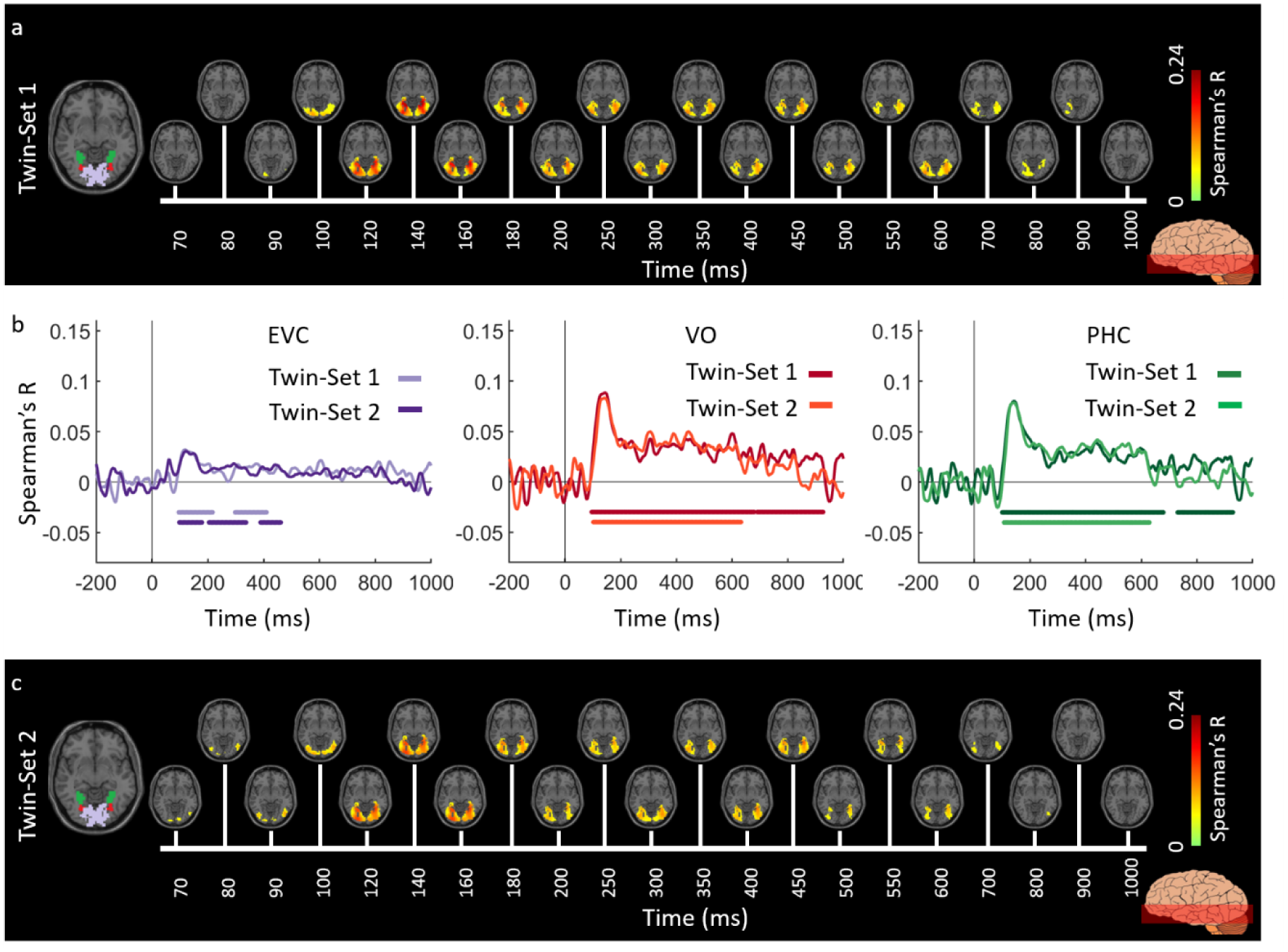
Experiment 1 Twins-set spatiotemporal neural dynamics of vision in ventral stream: (**a**) An axial slice encompassing early visual cortex (EVC) and ventral ROIs, ventral occipital (VO) area and parahippocampal cortex (PHC). The significant correlation maps for Twin-set 1 in this axial slice depicted over time (n = 15, cluster-definition threshold P < 0.001, cluster threshold P < 0.01); (**b**) The correlation time series are computed based on spatially restricted searchlight voxel-wise fusion analysis (see Methods). The depicted curves from left to right compare these time series in EVC and ventral ROIs VO and PHC for Twin-set 1 and Twin-set 2. Significant time points depicted with color coded lines below the graphs are determined with sign-permutation tests (n=15; P < 0.01 cluster-definition threshold, P < 0.01 cluster threshold); (**c**) The significant correlation maps for Twin-set 2 in the same axial slice as (a) depicted over time (n = 15, cluster-definition threshold P < 0.001, cluster threshold P < 0.01).

**Figure 3.**
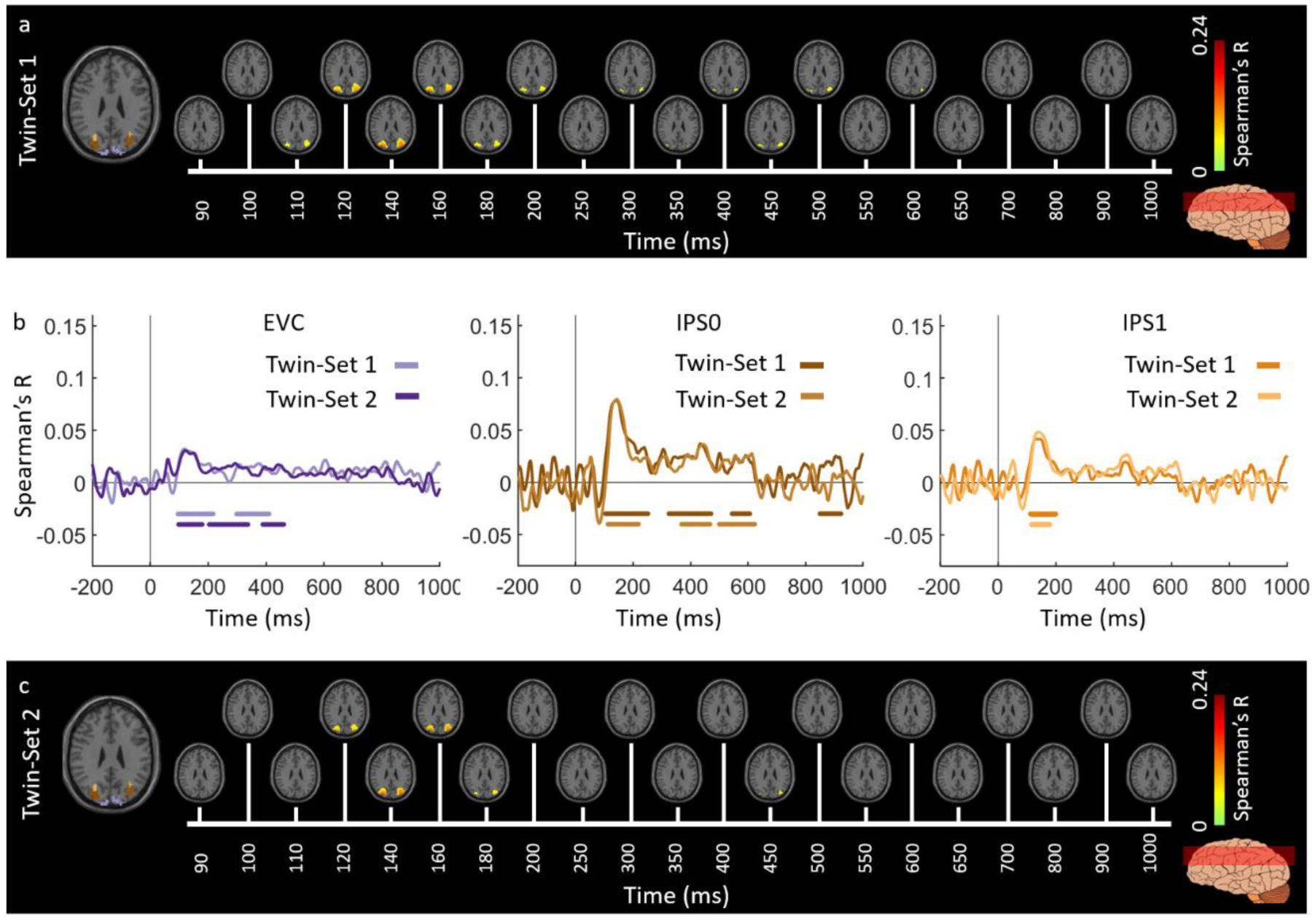
Experiment 1 Twins-set spatiotemporal neural dynamics of vision in dorsal stream: (**a**) An axial slice encompassing early visual cortex (EVC) and dorsal ROIs, inferior parietal sulcus (IPS0 and IPS1). The significant correlation maps for Twin-set 1 in this axial slice depicted over time (n = 15, cluster-definition threshold P < 0.001, cluster threshold P < 0.01); (**b**) The correlation time series are computed based on spatially restricted searchlight voxel-wise fusion analysis (see Methods). The depicted curves from left to right compare these time series in EVC and dorsal ROIs, IPS0 and IPS1 for Twin-set1 and Twin-set2. Significant time points depicted with color coded lines below the graphs are determined with sign-permutation tests (n=15; P < 0.01 cluster-definition threshold, P < 0.01 cluster threshold); (**c**) The significant correlation maps for Twin-set 2 in the same axial slice as (a) depicted over time (n = 15, cluster-definition threshold P < 0.001, cluster threshold P < 0.01).

To quantitatively compare the spatiotemporal fusion maps between the Twin-sets we performed spatially restricted voxel-wise fMRI-MEG fusion on five regions-of-interest (ROIs) (early visual cortex (EVC), ventral occipital area (VO), parahippocampal cortex (PHC), and inferior parietal sulci (IPS0 and IPS1). We averaged the correlation values over the voxels within each ROI resulting in one correlation time series per individual. Figures 2b and 3b compare the ROI-specific time courses for Twin-set 1 and 2. We observe that the two sets result in similar temporal dynamics within the regions of interest across ventral and dorsal pathways (All statistical tests are performed using permutation tests with cluster defining threshold P<0.01, and corrected significance level P<0.01; n=15). This demonstrates the reliability of the fusion method in reproducing similar patterns across similar visual experiences.

Next, we performed an ROI-based fMRI-MEG fusion following the method described in [8,9] (see Figure 4a). Studied ROIs (see details in Appendix D) include EVC, ventral regions (VO, TO, PHC) and dorsal regions (IPS0, IPS1, IPS2, and IPS3). As depicted in Figure panels 4bcde, the ROI-based fMRI-MEG fusion results in visually similar time series. Comparison of peak latency times show a peak of response around 120 msec in EVC and significantly later (two-sided hypothesis test, all P<0.01, FDR corrected), around 140 msec in ventral and dorsal ROIs. The peak latency and onset times with their corresponding 95% confidence intervals are reported in Table 1.

**Figure 4.**
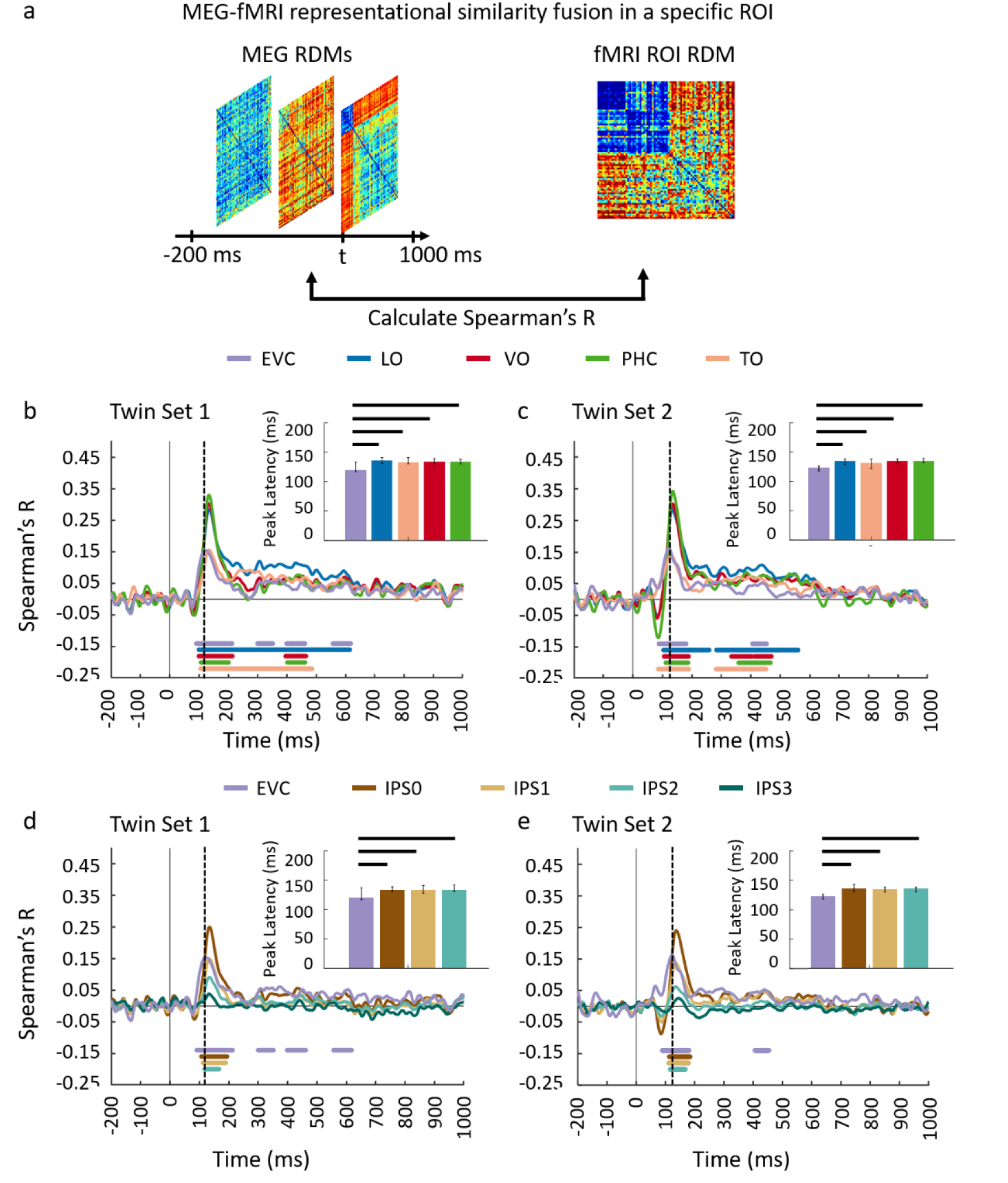
Comparing dorsal and ventral stream neural responses through similarity-based ROI fMRIMEG fusion. (**a**) The voxel patterns are extracted from each ROI to construct the fMRI ROI RDM. Then the ROI-specific fMRI RDM was compared with time-resolved MEG RDMs resulting in correlation time series for each region of interest; (**bc**) The fMRI-MEG fusion time series are depicted in EVC and ventral ROIs, LO, VO, PHC, and TO for Set 1 and 2, respectively; (**de**) The fMRI-MEG fusion time series are depicted in EVC and dorsal ROIs, IPS0-3 for Twin-set 1 and Twin-set 2, respectively. The color coded lines below the curves indicate significant time points (n = 15, cluster-definition threshold P < 0.01, cluster threshold P < 0.01) and the dashed vertical line in each plot indicates the peak latency in EVC time series. Peak latency times and their corresponding 95% confidence intervals for correlation time series in bcde are illustrated with barplots and error bars, respectively. Black lines above the bar indicate significant differences between conditions. 95% confidence intervals were found with bootstrap tests and barplots were evaluated with two-sided hypothesis tests; false discovery rate corrected at P<0.05.

**Table 1.**
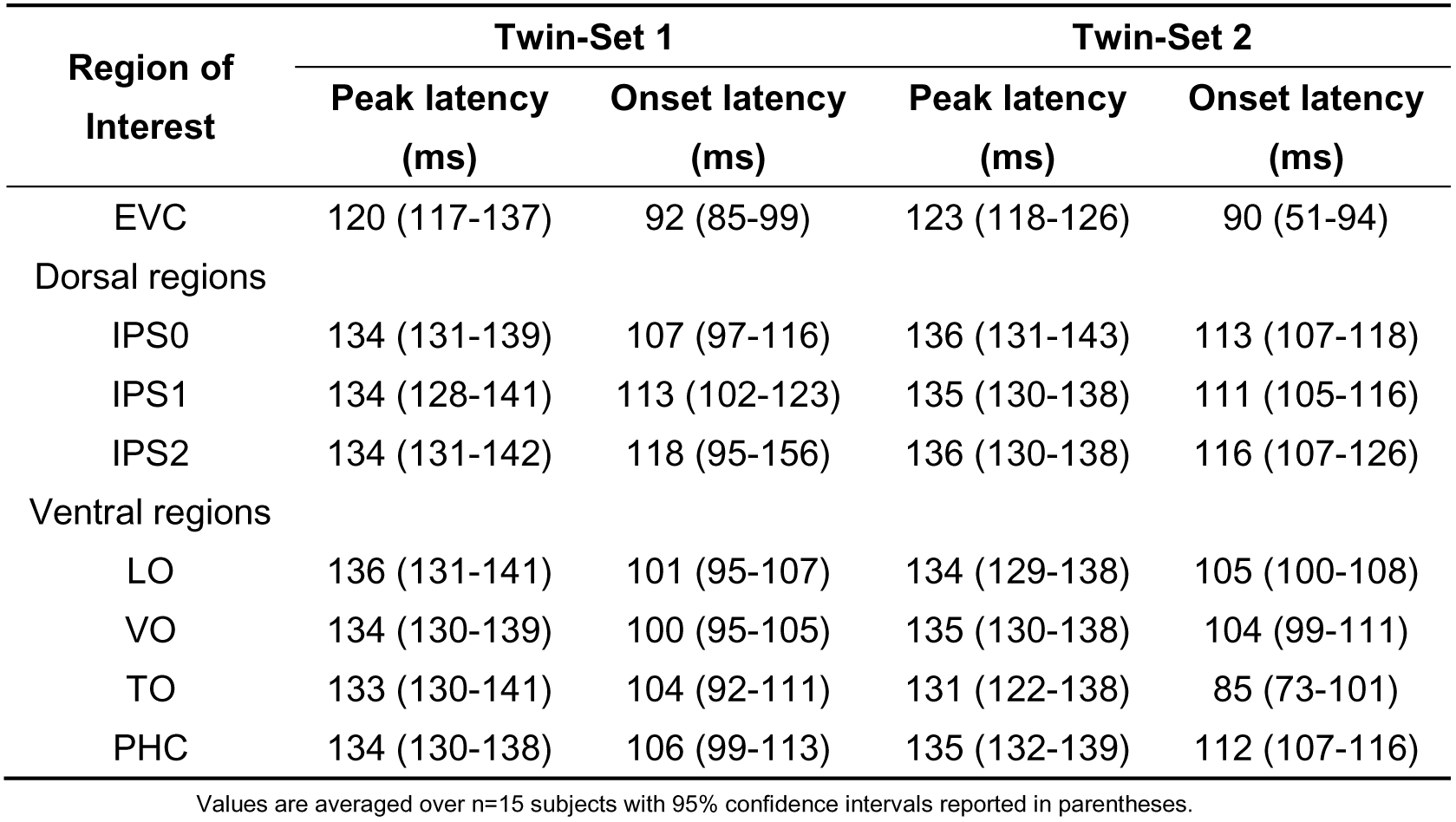
Peak and onset latency for fMRI-MEG fusion time series in EVC, ventral, and dorsal regions for Twin-Set 1 and Twin-Set 2 (Experiment 1).

### 3.2. Generalizability of similarity-based fMRI-MEG fusion method

We evaluated the generalizability of the fMRI-MEG similarity-based data fusion method by comparing results across participant groups presented with different images (Twins-Set and ImageNet-Set from Experiment 2 of [9]). We applied the fMRI-MEG similarity-based data fusion method to Twins-set and ImageNet-set, separately. Figures 5 and 6 display the spatiotemporal dynamics of visual perception for the two datasets along the ventral and dorsal visual pathways, respectively, over the first 1000 msec from stimulus onset. In both cases, the significant signals emerge in EVC around 70-80 msec after stimulus onset and and then in the anterior direction along the ventral stream (Figure 5ac), and across the dorsal stream up to the inferior parietal cortex (Figure 6ac). We determined significant spatiotemporal correlations with sign-permutation tests (n=15; P < 0.01 cluster-definition threshold, P < 0.001 cluster threshold).

**Figure 5.**
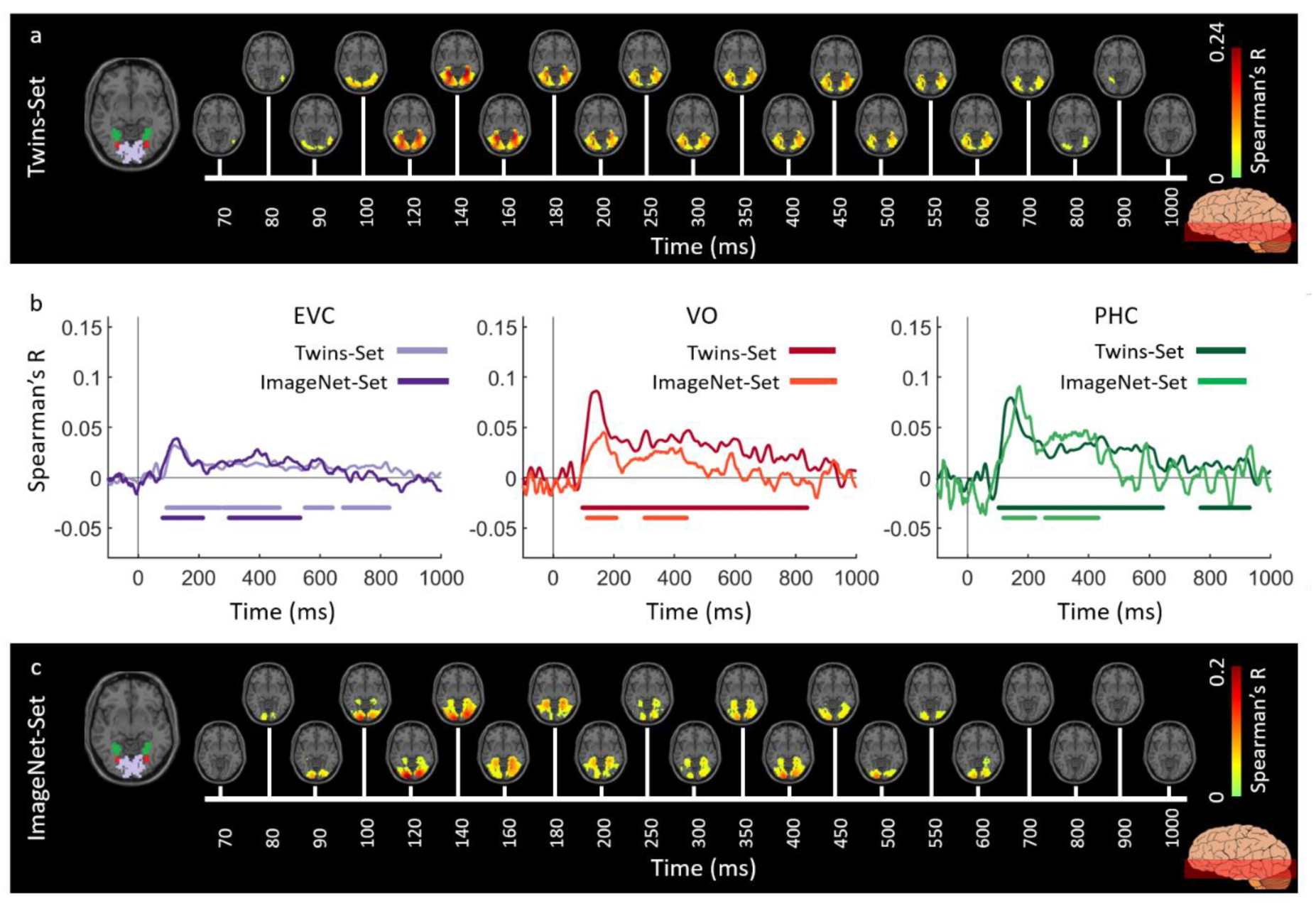
Spatiotemporal neural dynamics of vision in ventral stream: (**a**) An axial slice encompassing early visual cortex (EVC) and ventral ROIs, occipital (VO) area and parahippocampal cortex (PHC). The significant correlation maps for Twins-set (Experiment 1) in this axial slice depicted over time (n = 15, cluster-definition threshold P < 0.001, cluster threshold P < 0.01); (**b**) The correlation time series are computed based on spatially restricted searchlight voxel-wise fusion analysis (see Methods). The depicted curves from left to right compare these time series in EVC and ventral ROIs VO and PHC for Twins-set (Experiment 1) and ImageNet-set (Experiment 2). Significant time points depicted with color coded lines below the graphs are determined with sign-permutation tests (n=15; P < 0.01 cluster-definition threshold, P < 0.01 cluster threshold); (**c**) The significant correlation maps for ImageNet-set (Experiment 2) in the same axial slice as (a) depicted over time (n = 15, cluster-definition threshold P < 0.001, cluster threshold P < 0.01).

**Figure 6.**
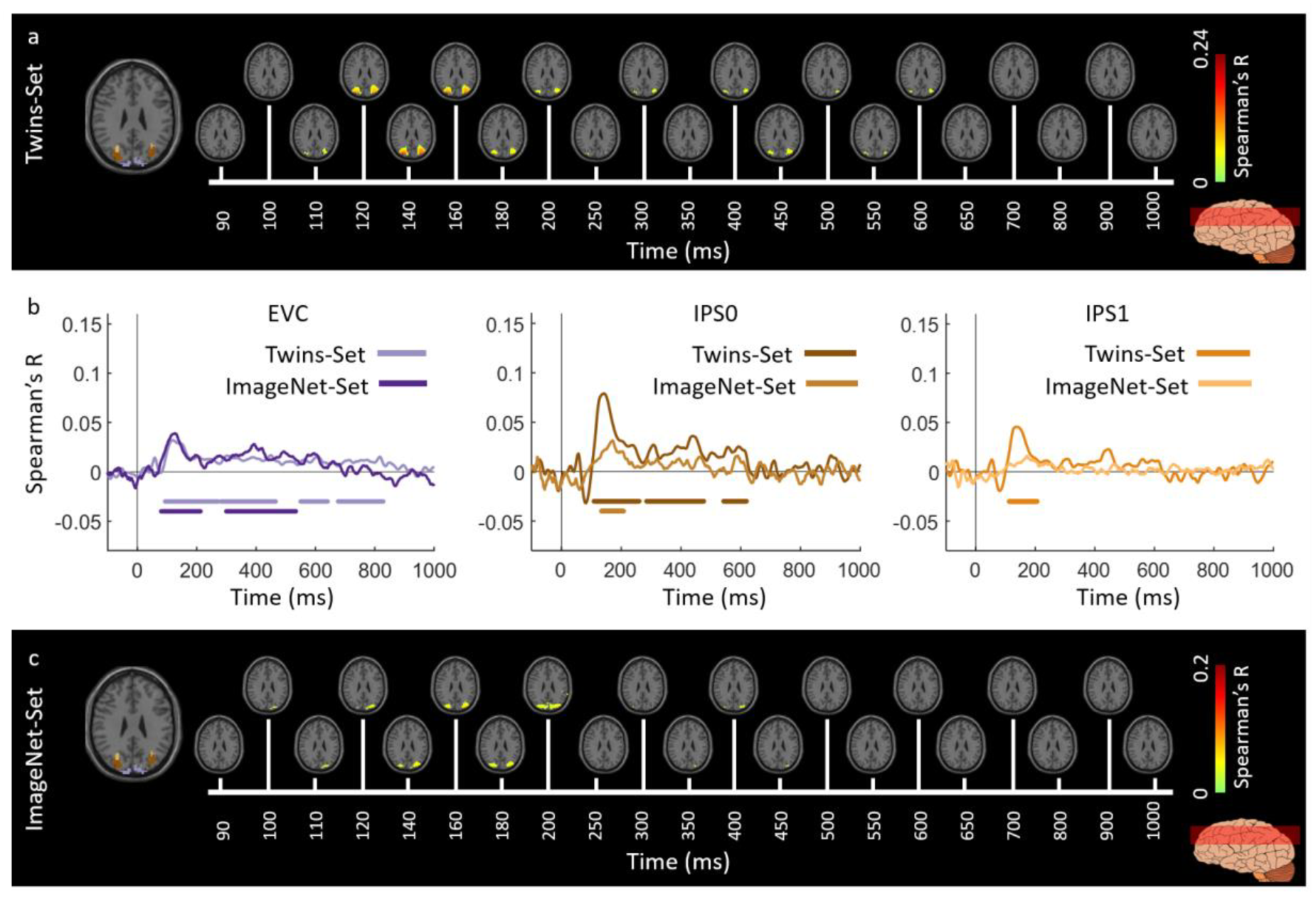
Spatiotemporal neural dynamics of vision in dorsal stream: (**a**) An axial slice encompassing early visual cortex (EVC) and dorsal ROIs, inferior parietal sulcus (IPS0 and IPS1). The significant correlation maps for Twins-set (Experiment 1) in this axial slice depicted over time (n = 15, cluster-definition threshold P < 0.001, cluster threshold P < 0.01); (**b**) The correlation time series are computed based on spatially restricted searchlight voxel-wise fusion analysis (see Methods). The depicted curves from left to right compare these time series in EVC and dorsal ROIs, IPS0 and IPS1 for Twins-set (Experiment 1) and ImageNet-set (Experiment 2). Significant time points depicted with color coded lines below the graphs are determined with sign-permutation tests (n=15; P < 0.01 cluster-definition threshold, P < 0.01 cluster threshold); (**c**) The significant correlation maps for ImageNet-set (Experiment 2) in the same axial slice as (a) depicted over time (n = 15, cluster-definition threshold P < 0.001, cluster threshold P < 0.01).

Figures 5b and 6b show qualitatively similar patterns of spatiotemporal fusion maps between the two datasets (Twins-Set and ImageNet-Set) by averaging the correlations over the voxels (spatially restricted voxel-wise fMRI-MEG fusion) in EVC, ventral regions (VO and PHC) and dorsal regions (IPS0 and IPS1).

To further investigate the similarities and dissimilarities between these two sets of data quantitatively, we performed ROI-based analyses. For each dataset, we correlated ROI-specific fMRI RDMs with time-resolved MEG RDMs resulting in correlation time courses shown in Figure 7abcd. We determined significant time points illustrated with color coded lines below the graphs with sign-permutation tests (n=15; P < 0.01 cluster-definition threshold, P < 0.01 cluster threshold). We observed that in both datasets, correlation time series peak significantly earlier in EVC compared to high level regions in ventral (VO and PHC) and dorsal (IPS0, IPS1, IPS2) pathways (two-sided hypothesis test, all P<0.01, FDR corrected) reflecting the established structure of the visual hierarchy. Peak and onset latencies of curves in Figure 7abcd and their corresponding 95% confidence intervals are reported in Table 2.

**Figure 7.**
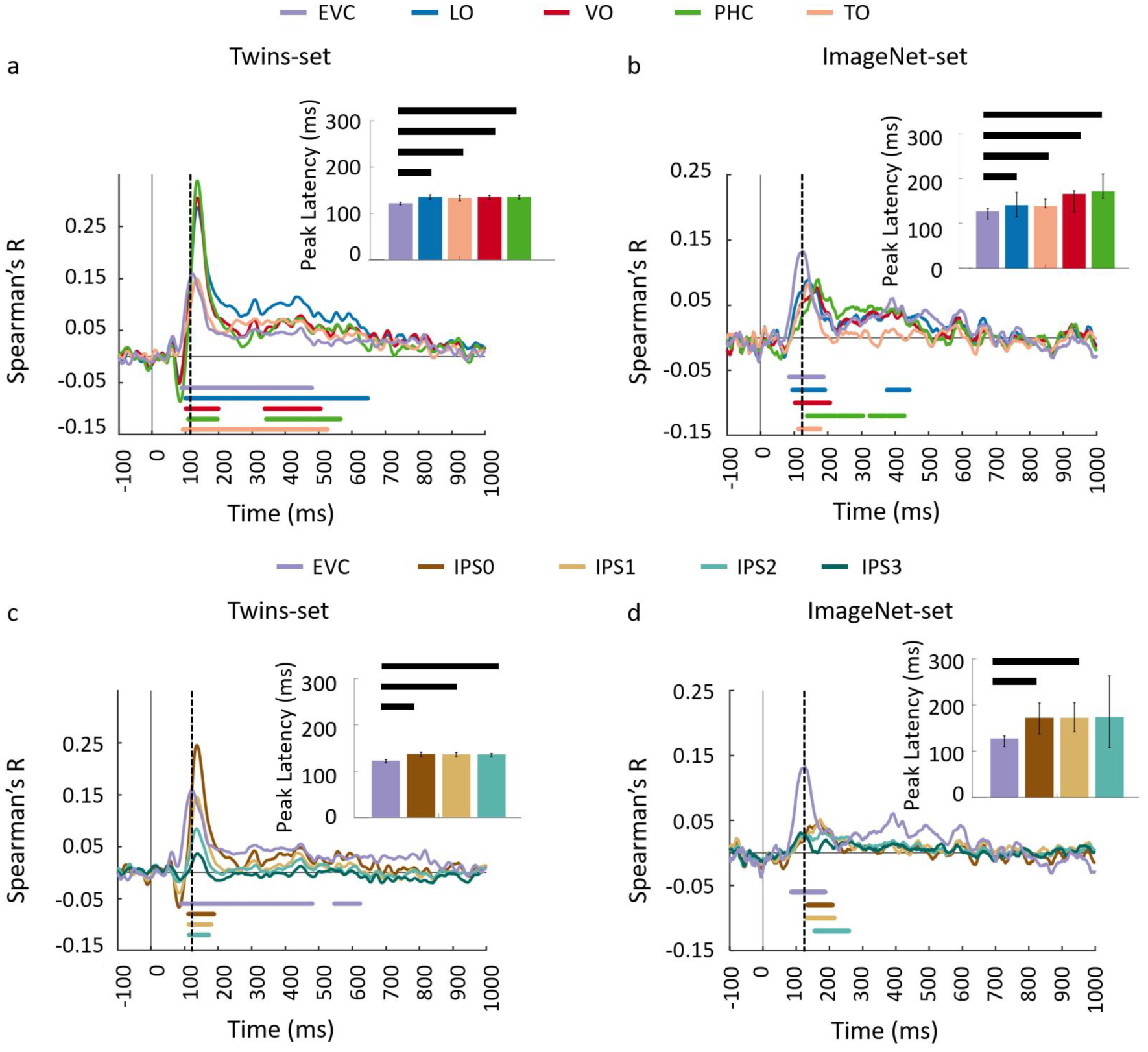
Comparing dorsal and ventral stream neural responses through similarity-based ROI fMRI-MEG fusion. The voxel patterns are extracted from each ROI to construct the fMRI ROI RDM. Then the ROI-specific fMRI RDM was compared with time-resolved MEG RDMs resulting in correlation time series for each region of interest; (**ab**) The fMRI-MEG fusion time series are depicted in EVC and ventral ROIs, LO, VO, PHC, and TO for Twins-set and ImageNet-set, respectively; (**cd**) The fMRI-MEG fusion time series are depicted in EVC and dorsal ROIs, IPS0-3 for Twins-set and ImageNet-set, respectively. The color coded lines below the curves indicate significant time points (n = 15, cluster-definition threshold P < 0.01, cluster threshold P < 0.01) and the dashed vertical line in each plot indicates the peak latency in EVC time series. Peak latency times and their corresponding 95% confidence intervals for correlation time series in bcde are illustrated with barplots and error bars, respectively. Black lines above the bar indicate significant differences between conditions. 95% confidence intervals were found with bootstrap tests and barplots were evaluated with two-sided hypothesis tests; false discovery rate corrected at P<0.05.

**Table 2.** Peak and onset latency for fMRI-MEG fusion time series in EVC, ventral, and dorsal regions for Twins-Set (Experiment 1) and ImageNet-Set (Experiment 2).

| Region of Interest | Twins-Set |  | ImageNet-Set |  |
| --- | --- | --- | --- | --- |
|  | Peak latency (ms) | Onset latency (ms) | Peak latency (ms) | Onset latency (ms) |
| EVC | 122 (119-125) | 89 (56-95) | 127 (110-133) | 84 (71-91) |
| Dorsal regions |  |  |  |  |
| IPS0 | 137 (131-141) | 111 (102-117) | 172 (137-204) | 133 (99-156) |
| IPS1 | 136 (132-140) | 111 (103-119) | 172 (142-205) | 137 (99-158) |
| IPS2 | 136 (132-138) | 112 (100-119) | 174 (108-263) | 103 (90-186) |
| Ventral regions |  |  |  |  |
| LO | 136 (131-141) | 102 (97-107) | 141 (115-169) | 96 (87-118) |
| VO | 136 (131-139) | 102 (95-109) | 166 (125-173) | 103 (95-125) |
| TO | 134 (128-140) | 90 (77-100) | 139 (135-154) | 115 (95-126) |
| PHC | 136 (132-139) | 109 (102-115) | 172 (156-210) | 121 (99-159) |

Values are averaged over n=15 subjects with 95% confidence intervals reported in parentheses.

## 4. Discussion

One of the central tenants of science is reproducibility [40–42]. This is especially relevant in cognitive neuroscience, as the surge of new cutting-edge multivariate analysis tools has made it possible to address questions that were previously untestable [38,43]. Here, we have demonstrated the reproducibility of the fMRI-MEG fusion method through the application of two separate experiments, testing the reliability and generalizability of the technique.

The first analysis compared neural data within the same subject groups across Twin image sets, with equalized low-level visual features, and sharing highly similar semantic concepts (e.g. a giraffe for a giraffe, a flower for a flower). We observed that the analysis yielded *reliably* consistent spatiotemporal dynamics, with each subset showing responses first in the occipital pole, and then signals in the anterior direction, with similar path in both ventral and dorsal streams. The strong agreement between the full brain spatiotemporal maps of the Twin-sets suggests that the signals detected by the fMRI-MEG data fusion method is likely to reliably reflect the representations of the stimulus. This sequence of representational signals followed the established spatial [44,45] and temporal [19,46] patterns associated with hierarchical visual processing.

This high degree of reliability, within subjects, across these separate but matched image-sets has pragmatic consequences in reinforcing the power and confidence of this method. The agreement within results of unimodal neuro-imaging have been shown have low test-retest reliability [28]. This realization has brought about a crisis of confidence in the replicability and reliability of published research findings within neuroscience [47,48]. Thus to mitigate these concerns the generation of new research discoveries requires that findings move beyond drawing conclusions solely on the basis of a single study. As such, we sought to extend this replication by comparing the newly collected Twins-Set data to previously published independent fMRI-MEG fusion data [9]. This analysis focused on the generalizability of this method across subject groups and to a wider range of stimuli.

Across the two experiments, findings from the full brain correlation maps and spatially restricted ROI analysis revealed that the fMRI-MEG data fusion captures the spatiotemporal dynamic patterns common to visual processing in a manner that *generalizes* across subjects and natural images. The results of both experiments illustrate the spatiotemporal progression associated with hierarchical visual processing [6,49,50], with significant signals emerging first in early visual areas and then along the ventral and dorsal pathways.

We found that overall this generalized response had similar expected spatiotemporal response patterns, but with greater variability in high-level visual areas. Previous work examining the replicability of neuroimaging data has shown that low-level brain functions (motor and sensory tasks), show generally less variance than high level cognitive tasks [51]. Moreover, between subject comparisons show significantly higher variability than within due to the increase sources of noise [52].

Cognitive neuroscientists have many methodological options for studying how experimental variables are systematically related to the spatiotemporal dynamics underlying a cortical process of interest. To date, much of our understanding has been advanced by methods that yield high spatial or temporal resolution, but not both. Separately, the limitations of these modalities in their application to understanding cognition are well known. Here we show that combining these methods through representational similarity analysis provides a reliable and generalizable tool for studying visual perception. This confirmatory replication within and between image-sets and subject-groups provides a spatiotemporal dynamic map of visual processing that can act as a guideline for further questions regarding the interactions between visual perception and cognitive factors, such as attention and memory.

## Author Contributions

conceptualization, Y.M., C.M., R.C., A.O.; investigation, Y.M., C.M., R.C., A.O.; methodology, Y.M., C.M., R.C., A.O.; data analysis, B.L., Y.M.; writing and editing, Y.M., C.M., R.C., A.O.; visualization, Y.M, C.M.; supervision, R.C., A.O.; funding acquisition, R.C., A.O.

## Acknowledgments

The experiments were conducted at the Athinoula A. Martinos Imaging Center at the McGovern Institute for Brain Research, Massachusetts Institute of Technology. The authors would like to thank Dimitrios Pantazis for helpful discussions. The data will be available online.

## Appendix A

*Experimental Design*: In both experiments, images were presented at the center of the screen for 500 msec with 6 and 4 degrees of visual angle in Experiment 1 and 2, respectively. The task was the same for both experiments: the images were overlaid with a black fixation cross at the center and participants performed an orthogonal vigilance task. The data were collected over one MEG session and two fMRI sessions in both experiments.

MEG session included 25 runs for Experiment 1 and 15 runs for Experiment 2. Images were presented once per run in randomized order with 1-1.2 sec stimulus onset asynchrony (SOA) in Experiment 1 and twice per run with 0.9-1 sec OSA in Experiment 2. The participants were instructed to press a button and make an eye blink upon detection of a specific image (depicting an eye in Experiment 1 and a paper clip in Experiment 2) presented every 3-5 trials randomly.

The two fMRI sessions consisted of 11-15 runs in Experiment 1 and 9-11 runs in Experiment 2. Images were presented once per run in random order. In both studies 39 null trials were randomly distributed in each run. During the null trials the fixation cross changed color for 100 msec and subjects were instructed to respond to this color change by pressing a button. The trial duration was 3 sec in both studies. Images were presented for 500 msec and stimulus onset asynchrony was 3 sec, or 6 sec (if a preceding null trial).

## Appendix B

### MEG Acquisition and Analysis

MEG data were acquired from a 306-channel Elekta neuromag TRIUX system with sampling rate of 1 kHz and filtered by a 0.03 to 330 Hz band-pass filter. Participants’ head position was measured prior and during the recording with 5 coils attached to their head. A maxfilter was applied for temporal source space separation and correcting for head movements [53,54]. Data were analyzed using Brainstorm software [55] to extract trials from −200 msec to 1000 msec with respect to image onset in Experiment 1 and from −100 msec to 1000 msec relative to image onset in Experiment 2. The mean of the baseline was removed and the data were smoothed with a 30 Hz low-pass filter. For each condition and participant, we obtained 25 trials in Experiment 1 and 30 trials in Experiment 2.

Multivariate pattern analysis was used to determine the similarity relations between visual conditions (Figure 1b). At each time point t, MEG sensor data were arranged in pattern vectors resulting in N vectors per condition at that time point with N being the number of trials per condition. Then, for each pair of conditions a support vector machine (SVM) classifier was trained to discriminate the conditions based on the MEG pattern vectors at each time point. The performance of the classifier computed with leave-one-out cross validation procedure was considered as a measure of dissimilarity between those pairs of conditions and used to construct a C x C (C=156 in Experiment 1 and C=118 in Experiment 2) representational dissimilarity matrix (RDM) at each time point. Each row and column in this matrix are indexed by a specific condition (image) and each matrix entry shows how dissimilar the corresponding conditions are based on the MEG signals at a specific time point. These matrices are symmetric with an undefined diagonal. To overcome computational complexity and reduce noise, trials were randomly sub-averaged in groups of 3 in Experiment 1, and groups of 5 in Experiment 2, before entering them into multivariate analysis.

### fMRI Acquisition and Analysis

The MRI data of both experiments were acquired with a 3T Trio Siemens scanner using a 32-channel head coil. The structural images were acquired using a standard T1-weighted sequence (192 sagittal slices, FOV = 256 mm2, TR = 1900 msec, TE = 2.52 msec, flip angle = 9°).

Functional data of Experiment 1 was acquired over 11-15 runs of 305 volumes in two sessions with gradient-echo EPI sequence: TR =2000 msec, TE = 29 msec, flip angle = 90°, FOV read = 200 mm, FOV phase = 100%, bandwidth 2368 Hz/Px, resolution = 3 mm3, slice gap 20%, slices = 33, ascending interleaved acquisition). Functional data of Experiment 2 was acquired over 9-11 runs of 648 volumes in two sessions with gradient-echo EPI sequence: TR =750 msec, TE = 30 msec, flip angle = 61°, FOV read = 192 mm, FOV phase = 100% with a partial fraction of 6/8, through-plane acceleration factor 3, bandwidth 1816 Hz/Px, resolution = 3 mm3, slice gap 20%, slices = 33, ascending interleaved acquisition).

The fMRI data in both studies were preprocessed using SPM software. For each participant, fMRI data were slice-time corrected, realigned and co-registered to the T1 structural scan of the first session, and finally normalized to the standard MNI space. We applied a general linear model (GLM) to estimate the fMRI responses to the 156 and 118 image conditions in Experiment 1 and 2, respectively. Stimulus onsets and durations, as well as motion and run regressors were included in the GLM. All the regressors were convolved with a hemodynamic response function (canonical HRF) with time resolution of TR/N and onset of the first slice (TR = 2000 msec, N = 33 slices). By contrasting each condition against the implicit baseline, we converted the condition-specific GLM estimates into t-value maps (resulting in 156 condition-specific t-maps for Experiment 1 and 118 condition-specific t-maps for Experiment 2).

We then applied the searchlight analysis method to create the fMRI RDMs [56,57] for each subject, separately. At each voxel *v*, condition-specific t-value pattern vectors were extracted in a sphere centered at voxel *v* with a 4-voxel radius. Dissimilarities between these pattern vectors were computed in a pairwise manner (1-Pearson’s R). Then these dissimilarities were entered into an RDM matrix in which its rows and columns were indexed by the image conditions. This process resulted in a 156 x 156 RDM at each voxel for fMRI data of each subject in Experiment 1 and a 118 x 118 RDM at each voxel for fMRI data of each subject in Experiment 2.

## Appendix C

*Full Brain fMRI-MEG Fusion:* The assumption in the fusion analysis is that the pairwise condition-specific relations (in the form of RDM representations) are preserved across MEG and fMRI data patterns. For example, if two images result in similar patterns in MEG data, they produce similar patterns in fMRI data as well. Comparison of pairwise similarity relations across MEG and fMRI makes it possible to relate representations at specific time points (in MEG) with specific locations (in fMRI). For both datasets, we performed within subject analysis comparing averaged MEG RDMs over the 15 subjects, with subject-specific fMRI RDMs. We computed the correlations (Spearman’s R) between MEG RDMs at each time point and fMRI RDMs at each voxel. These computations resulted in a 3D correlation map over the whole brain at each time point. Repeating the analysis over time yielded a spatiotemporally resolved view of visual information in the brain.

## Appendix D

### Region-of-Interest Analysis

*Spatially Restricted Searchlight Voxel-wise fMRI-MEG Fusion*: This analysis followed the regular voxel-wise full brain fMRI-MEG fusion but in spatially restricted regions of the brain. In detail, the search-light method was performed within the ROI and an RDM matrix was created at each specific voxel by comparing condition-specific fMRI pattern responses within the searchlight sphere centered at the voxel (1-Pearson’s R). Then the time-resolved MEG RDMs were compared with these fMRI RDMs (Spearman’s R) resulting in correlation time series within the ROI which were then averaged resulting in one correlation time-course per ROI and per subject. This ROI analysis method is directly comparable with the full brain fusion analysis and it is a convenient way to quantitatively compare the spatiotemporal dynamics of fusion movies within specific brain regions.

*ROI-based fMRI-MEG Fusion*: This analysis method followed [8,9]. We extracted the voxel patterns within each ROI and then computed the condition-specific pairwise dissimilarities (1-Pearson’s R) to create a single RDM matrix per ROI. Thus we obtained one fMRI RDM for each ROI per subject. The fMRI ROI RDMs were averaged over subjects and then compared with subject-specific MEG RDMs over time by computing their Spearman’s R correlations. This process results in time-course of MEG-fMRI fusion in each ROI per subject.

Specifically, we extracted the following ROIs: one early visual cortex (combining V1, V2, V3), four ventral visual areas (lateral occipital (LO1&2), temporal occipital (TO), ventral occipital (VO1&2), and parahippocampal cortex (PHC1&2) and four parietal areas (intraparietal cortex including IPS0, IPS1, IPS2, and IPS3). The ROI definition followed [9] based on the probabilistic maps of [58].

## Appendix E

### Statistical Testing

For statistical inference, we used non-parametric statistical tests [59,60] assessed by permutation-based cluster-size inference. The null hypothesis was zero for MEG-fMRI correlation time series and 4D spatiotemporal correlation maps. In all cases, we used 1000 permutations, 0.01 cluster defining threshold and 0.01 cluster threshold for time series; and 0.001 cluster defining threshold and 0.01 cluster threshold for 4D maps.

For peak and onset latency of the time series statistical assessments, bootstrap tests were performed. We bootstrapped the subject-specific time series for 1000 times to estimate an empirical distribution over peak and onset latency of the subject-averaged time courses and use it to define 95% confidence intervals. For peak to peak latency comparisons, 1000 bootstrapped samples of two peak differences were obtained. The null hypothesis was rejected if the 95% confidence interval of the peak latency differences did not include zero.

